# Birds that don’t exist: niche pre-emption as a constraint on morphological evolution in the Passeroidea

**DOI:** 10.1101/2025.09.24.678290

**Authors:** Stephanie Y. Chia, Anshuman Swain, Nathaniel Josephs, Lizhen Lin, William F. Fagan

## Abstract

Understanding why some viable body forms never evolve can reveal how ecological and evolutionary forces shape biodiversity. We investigate this question in the Passeroidea, a large group of songbirds, by analyzing their morphological trait space using topological data analysis and ancestral state reconstruction. We identify a long-standing morphological gap densely surrounded by extant species but unoccupied throughout passeroid diversification. The gap patterns deviate from neutral expectations and show no evidence of past occupation, rendering neutral evolution and extinction unlikely. Similar morphologies exist in other bird lineages, ruling out intrinsic constraints or niche absence. Geographic distributions and traits of passeroids versus non-passeroid gap occupants point to competitive exclusion as the plausible explanation: early-colonizing territorial specialists outside the Passeroidea may have preemptively occupied key habitats, limiting evolutionary opportunities for later-arriving lineages. We demonstrate how historical contingency can shape macroevolutionary outcomes, and introduce a generalizable framework for investigating structural gaps in trait evolution.

## 1. Introduction

Birds, with over ten thousand species, exhibit remarkable morphological diversity. This variation in body form underlies their ecological success, enabling species to exploit a wide range of environments and resources. Morphological traits are tightly linked to ecological function. For example, beak shapes reflect feeding specialization such as seed-cracking or insect-catching (Herrel *et al*. 2005; Lack 1983), wing and tail morphologies influence flight style and migratory ability (Norberg 1990), leg length relates to habitat use (from long-legged waders in aquatic habitats to short-legged aerial insectivores that minimize drag; Zeffer *et al*. 2003), and body mass affects thermoregulation, starvation resistance, and predator-prey dynamics (Peters 1986).

Together, these traits define a multidimensional morphological trait space, or **morphospace**, wherein each species occupies a unique position corresponding to its ecological niche (Hutchinson 1957; Pigot *et al*. 2020). Unoccupied regions in this morphospace, so-called **morphological gaps**, may reflect inviable trait combinations, ecological niches that do not exist or are biophysically or biogeographically inaccessible, or viable strategies that have not yet evolved. Identifying and understanding these gaps offers insight into the ecological and evolutionary forces that shape the limits of biodiversity.

Detecting such gaps in high-dimensional space is challenging, especially when these gaps lie within otherwise densely occupied regions. Such “holes” can be obscured by dimensionality reduction techniques commonly used in trait-based analysis. To address this, Blonder (2016) introduced a probabilistic framework to detect holes in high-dimensional hypervolumes by comparing observed trait distributions to convex or maximal expectations. This kernel density-based approach opened new avenues for exploring trait space, including studies of extinction risk in birds (Ali *et al*. 2023). However, it requires selecting a bandwidth and may be sensitive to uneven trait distributions, which are common for data in morphospaces.

As an alternative, we explore **persistent homology**, a method from topological data analysis that detects topological features in multidimensional point clouds (Edelsbrunner *et al*. 2002) without assuming smoothness or evenness in data distribution. It works by progressively connecting nearby species in the trait space and detecting loops that persist across a range of distance thresholds, thereby revealing robust gaps. Imagine a flooded landscape (morphospace) with many mountain peaks (species). As water level gradually drops (increasing distance threshold), peaks connect into ridges; when ridges form a loop, a lake appears (birth of a morphological gap). As water level continues to drop, the lake eventually disappears as the terrain fills in (death of the gap). The longer the lake persists across water levels, the more robust the gap is. Although the potential of persistent homology for studying ecological niche hypervolumes and trait distributions has been noted (Conceição & Morimoto 2022), it remains largely unexplored.

We apply persistent homology to examine the structure and temporal dynamics of the morphospace in the superfamily Passeroidea, a large clade of songbirds, as a case study. By combining persistent homology with ancestral state reconstruction, we detect and track the existence of morphological gaps through evolutionary time. Specifically, we ask: (1) Are there combinations of morphological traits that do not exist among extant passeroids? (2) If so, what ecological or evolutionary processes explain these absences?

To address these questions, we consider several hypotheses (**Fig. 1**). Morphological gaps may reflect intrinsic constraints, such as biomechanical, developmental, or genetic limitations that make certain trait combinations inviable, either lying outside of the observed range of extant species (**Hypothesis A**), or due to more subtle limitations within the realized morphspace (**Hypothesis B**). Gaps might also emerge as a neutral outcome of trait diversification, reflecting viable trait combinations that have not evolved simply due to chance or insufficient evolutionary time (**Hypothesis C**). Other possibilities include extinction of lineages that once occupied the gap (**Hypothesis D**), artifacts of missing data (**Hypothesis E**), or ecological mismatch whereby suitable niches for the morphology do not exist or are inaccessible (**Hypothesis F**). Lastly, a morphology might be viable and realized in other clades but absent from the focal clade due to competitive exclusion (**Hypothesis G**). We assess these seven possibilities by examining the evolutionary persistence of morphological gaps, their positions in morphospace, evidence of historical occupation by ancestral or extinct species, and the ecological and phylogenetic context of species that are morphologically similar.

**Figure 1.**
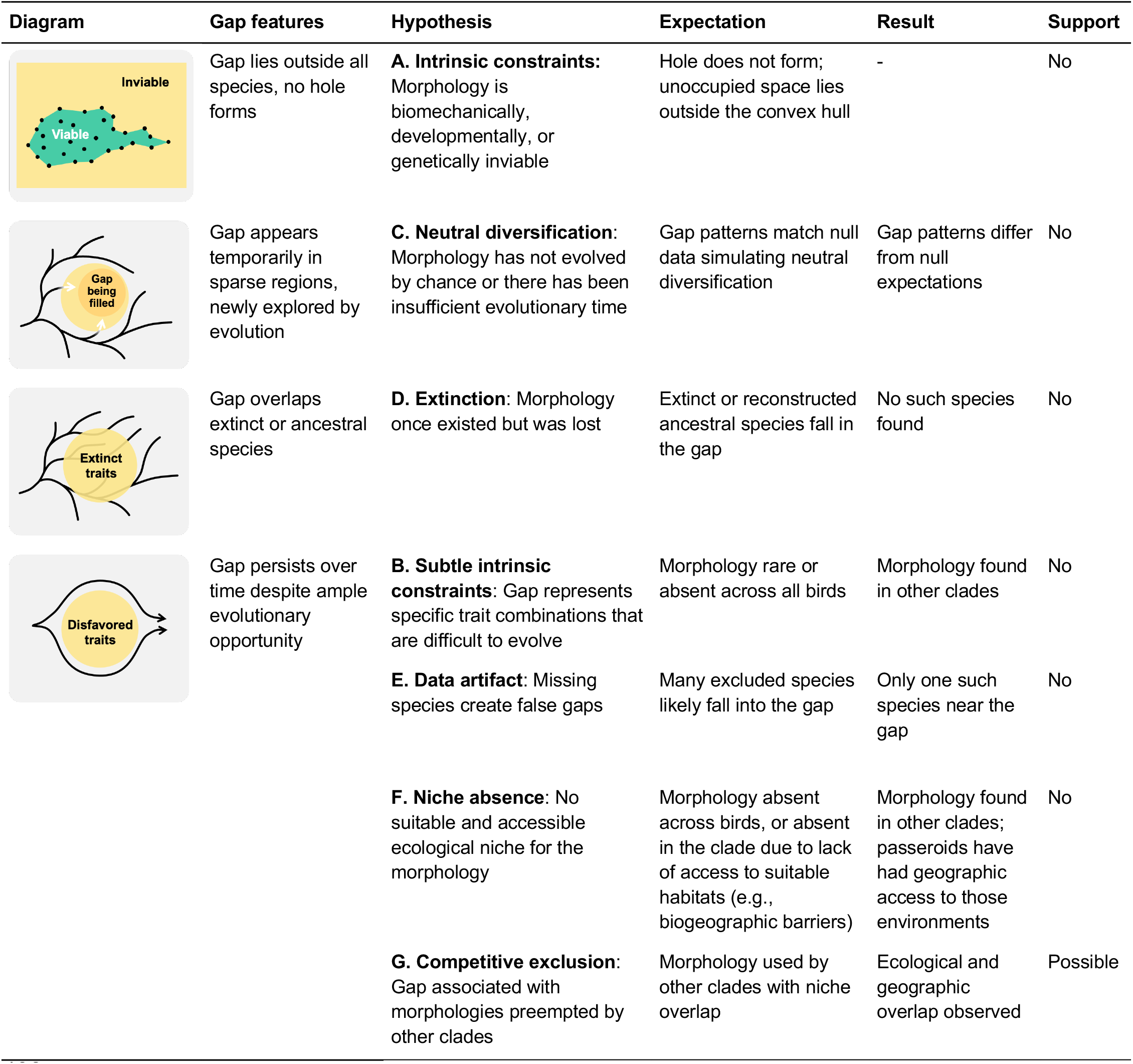
Conceptual framework and evaluation of hypotheses for the presence of a morphological gap in the Passeroidea trait space. Diagrams illustrate gap features associated with each hypothesis, with yellow area indicating unoccupied regions of morphospace and black curves showing evolutionary trajectories through morphospace. Hypotheses D through G share the same gap diagram. The expectation, observed result, and level of support for each hypothesis are summarized.

## 2. Materials and methods

### 2.1 Morphological and ecological traits

We analyze the superfamily Passeroidea as a case study to explore morphological macroevolution. This large monophyletic clade (Mackiewicz *et al*. 2019) within the passerines includes approximately 1,400 species, including sparrows, finches, tanagers, pipits, and many types of warblers. Members of the Passeroidea share relatively similar morphology, resulting in a densely populated morphospace that is well suited for detecting robust topological features.

We use ten morphological traits obtained from the AVONET dataset (Tobias *et al*. 2022): beak length from tip to skull along culmen, beak length from tip to anterior edge of nares, beak width, beak depth, tarsus length, wing length (carpal joint to wingtip), secondary length (carpal joint to tip of outermost secondary), tail length, hand-wing index, and body mass. We excluded 40 extant species that had at least one morphological trait (excluding body mass) referenced from a closely related species, resulting in a final dataset of 1,378 extant species. All traits were log-transformed (except the hand-wing index) and standardized to mean of zero and standard deviation of one. We applied principal component analysis (PCA) and retained the first four principal components (PCs), which together explained 95% of the total variance.

Morphological data for 60 recently extinct passeroid species (i.e., extinct within the past ∼130,000 years) were obtained from the AVOTREX dataset (Sayol *et al*. 2023). Secondary length was derived as the sum of wing length and Kipp’s distance, whereas the hand-wing index was calculated as Kipp’s distance divided by wing length. These extinct species were projected onto the morphospace defined by extant species’ PCs.

Trophic niche, primary lifestyle, and breeding range centroid and range size were also obtained from the AVONET dataset (Tobias *et al*. 2022). Territoriality was sourced from Tobias et al. (2016). BirdTree taxonomy (Jetz *et al*. 2012) was adopted to compile all trait data.

### 2.2 Ancestral state reconstruction and trait evolution simulations

Phylogenetic tree data were retrieved from BirdTree.org by randomly sampling 1,000 trees from Hackett backbone trees. A consensus tree was generated using majority rule topology and least squares edge length estimation.

Ancestral state reconstruction for each PC was performed under a Brownian Motion model using generalized least squares (**Fig. S1**). The resulting root values and evolutionary rate (s^2^) were used to generate ten null trait datasets under Brownian motion using the same consensus tree. Trait values for all datasets were linearly interpolated at 1-million-year intervals based on the tip (present-day trait values) and ancestral node values.

All phylogenetic analyses and simulations were performed in R using the package *phytools* (Revell 2024).

### 2.3 Gap identification and characterization

At each time slice, we applied persistent homology to detect morphological gaps in the four-dimensional morphospace defined by the first four PCs. Persistent homology is a topological data analysis tool that can identify features such as holes in multidimensional point clouds (Otter *et al*. 2017). We used Vietoris-Rips filtration based on Euclidean distance to detect loop-like structures (H1 features) which we interpret as morphological gaps. For each detected gap, we extracted the coordinates of the points forming the loop for downstream analysis. Computations of persistent homology were performed using the *Dionysus* C++ library via the R package *TDA* (Fasy *et al*. 2025).

Each gap’s **topological persistence** (i.e., the difference between “gap birth” and “gap death” values) was used to quantify the robustness of the structure across distance scales in the morphospace. We also computed the gap **centroid** as the mean location of its vertices, gap **size** as the mean distance from the centroid to its vertices, and **sparsity** as the mean distance from the centroid to the nearest 5% of species (69 of 1,378 data points), serving as an intuitive, distance-based measure of local density. All these metrics are in PC distance unit (dimensionless). To account for the irregular shapes of topological gaps, we defined a species as being **within** or **near** a gap based on scaled distances from the gap centroid, using half and full gap sizes as heuristic thresholds, respectively.

We extracted notable gap structures with topological persistence > 0.4 (PC unit) and linked gaps between consecutive time slices into **gap series** if the centroids were within 1 unit. For each gap series, we calculated **evolutionary lifespan** (duration of gap persistence in millions of years), average size, and average sparsity.

## 3. Results

### 3.1 A long-standing morphological gap in Passeroidea

Our analysis included 1,378 extant species (97.2%) in the superfamily Passeroidea after excluding 40 species with incomplete morphological data. We summarized ten morphological traits (available for all 1,378 species) using principal component analysis (PCA), with the first four principal components accounting for 95% of total variance. These axes primarily represent body size (PC1), beak thickness (PC2), hand-wing index (PC3), and beak length (PC4) (**Fig. 2, Table S1**).

**Figure 2.**
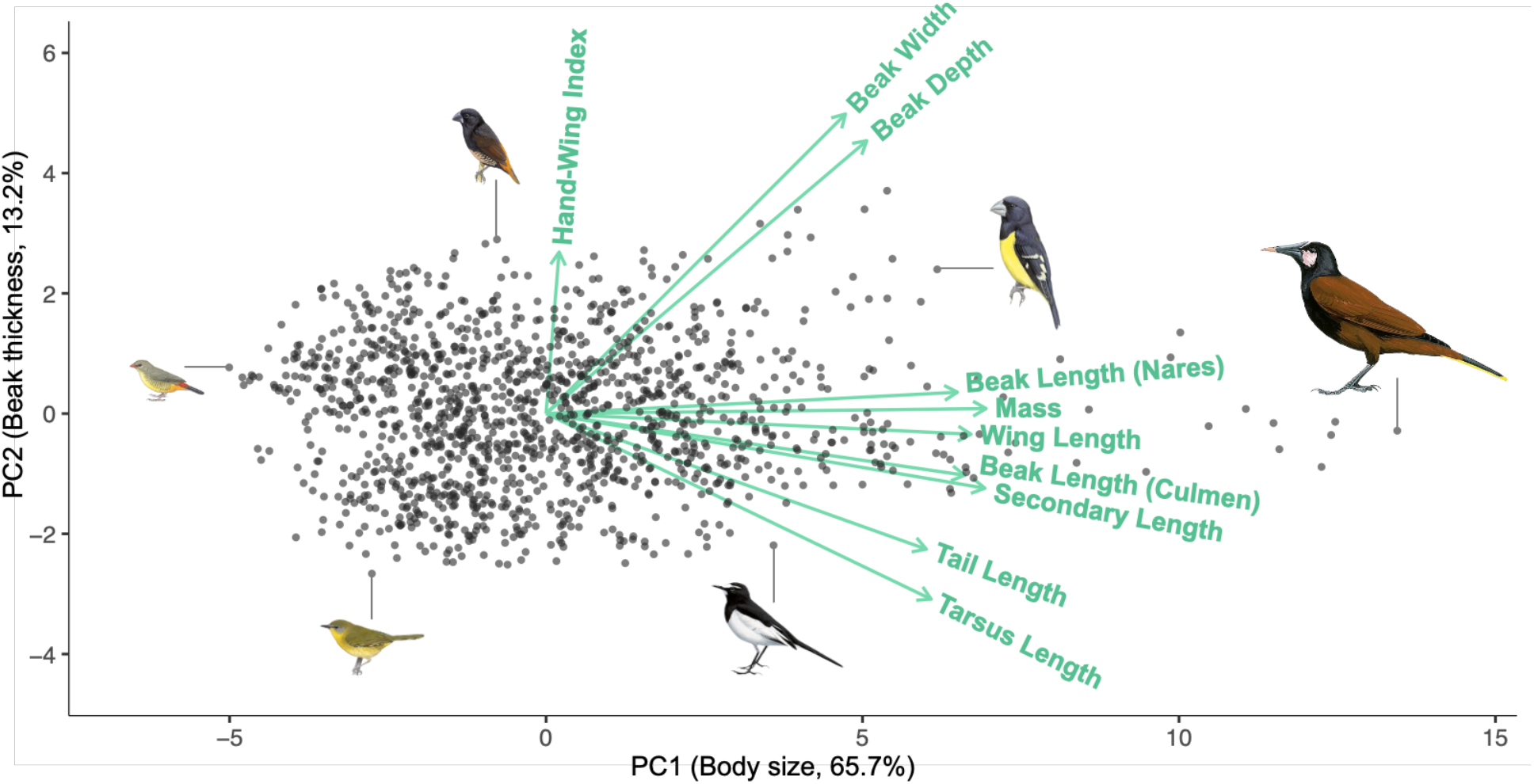
Morphological trait space of 1,378 extant Passeroidea species. Each point represents a species positioned by its scores on the first two principal components, which together explain 78.9% of the total morphological variance. PC1 primarily reflects body size, and PC2 corresponds to beak thickness. Green arrows indicate the loadings of the original morphological traits on the PC axes. Species illustrations (from left to right: Zebra Waxbill, Hooded Yellowthroat, Bismarck Munia, Japanese Wagtail, Spot-winged Grosbeak, and Baudo Oropendola) are from Birds of the World, Cornell Lab of Ornithology.

To identify morphological gaps, we applied persistent homology to the four-dimensional trait distributions with ancestral traits reconstructed at 1-million-year intervals. Within each time slice, we detected loop-like topological features representing unoccupied gaps in morphospace. Each gap was characterized by its **topological persistence**, a measure of robustness across distance scales in morphospace. We then linked recurring gaps across consecutive time slices to determine their **evolutionary lifespan** (in millions of years).

One particular morphological gap (hereafter the **focal gap**) stood out by persisting for 7 million years and exhibited high topological persistence within each time slice (**Fig. 3**; **Table S2**). This gap occurred in a densely occupied region of morphospace and corresponded morphologically to a medium-sized songbird with a relatively thin beak (**Figure S2**). Despite abundant surrounding species, this trait combination remained consistently unoccupied throughout evolutionary time in the Passeroidea.

**Figure 3.**
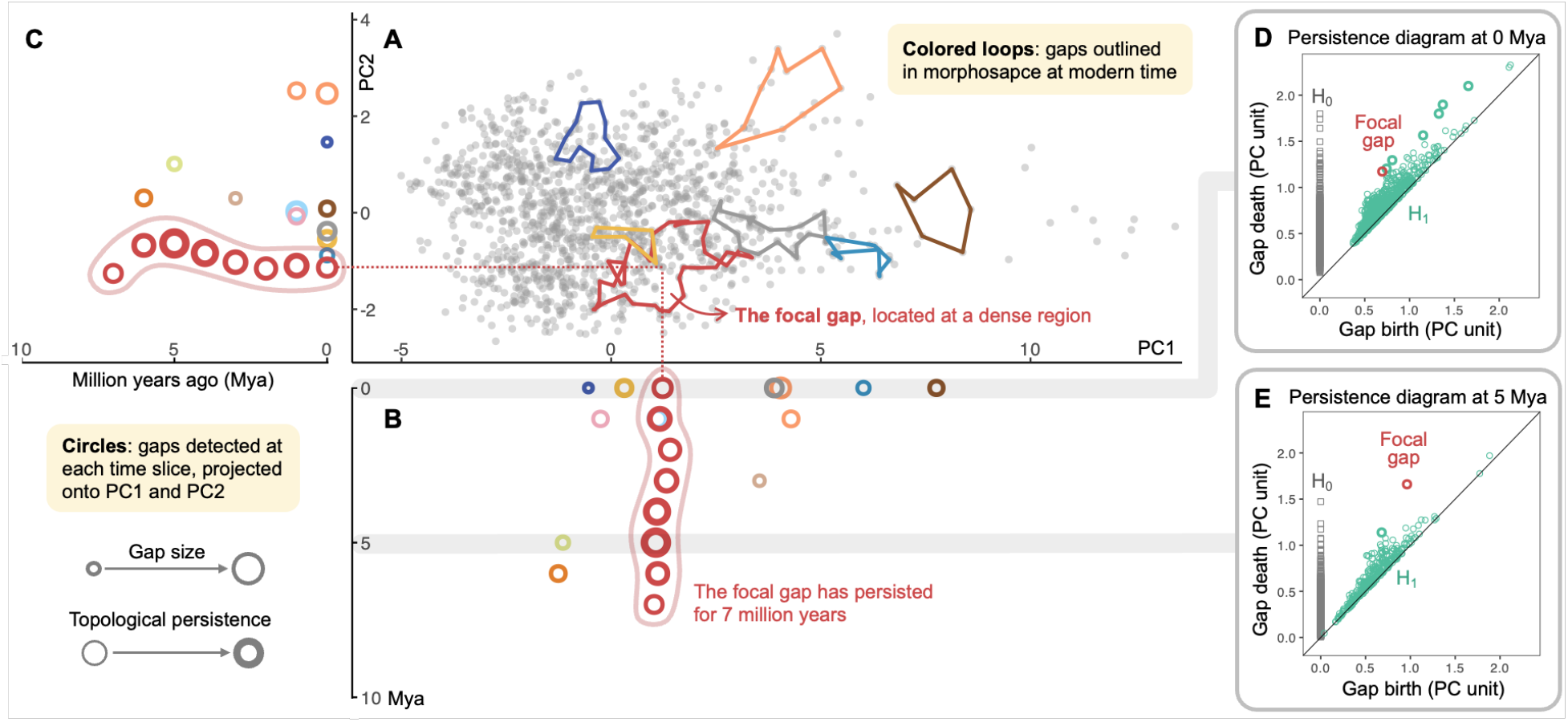
Temporal dynamics of identified gaps in Passeroidea morphospace. **A** shows extant passeroids in the morphospace formed by PC1 and PC2, with colored loops marking notable modern gaps. Note that this is a two-dimensional projection of a four-dimensional space; although some loops appear folded and many data points fall within the loops, they are actually unoccupied gaps in the full 4-D trait space. **B** and **C** indicate the position of detected 4-dimensional gaps projected onto PC1 and PC2, respectively, at each time slice (in million years ago, Mya). Each circle represents a gap, with its size reflecting relative gap size (mean distance from the gap centroid to its vertices), and stroke thickness indicating topological persistence (i.e., persistence of the gap structure across distance scales in trait space). Circles with the same colors depict gap series, representing gaps located in similar regions of morphospace across consecutive time slices. One particular gap (the focal gap) has persisted for 7 million years at a relatively dense region in the morphospace. Beyond the focal gap, only one other gap lasts more than 1 million years. **D** and **E** show the detected topological features of two homology groups, H_0_ (connected components; gray squares) and H_1_ (loops; green circles) at the present time and at 5 Mya, respectively. This study focuses on H_1_ features (gaps), with the focal gap shown in red throughout all panels.

### 3.2 The focal gap deviates from neutral diversification expectations

To assess whether the focal gap could result from a neutral diversification process, we generated 10 null trait datasets simulating Brownian motion evolution using the same phylogeny, root values, and evolutionary rates as in the empirical data.

Compared to gaps detected in the null datasets, the focal gap was notable in that it exhibited both a long evolutionary lifespan and high topological persistence within a densely occupied region of morphospace (**Fig. 4**). At the modern time slice, most gaps with high topological persistence in the null datasets were found in sparser regions of morphospace, whereas the focal gap was situated in a densely populated area (**Fig. 4A**). Across time, gap series in the null datasets were typically short-lived; only four persisted for more than 4 million years among all gap series detected across the ten null datasets, and none of these occurred in regions as densely occupied as the focal gap (**Fig. 4B**).

**Figure 4.**
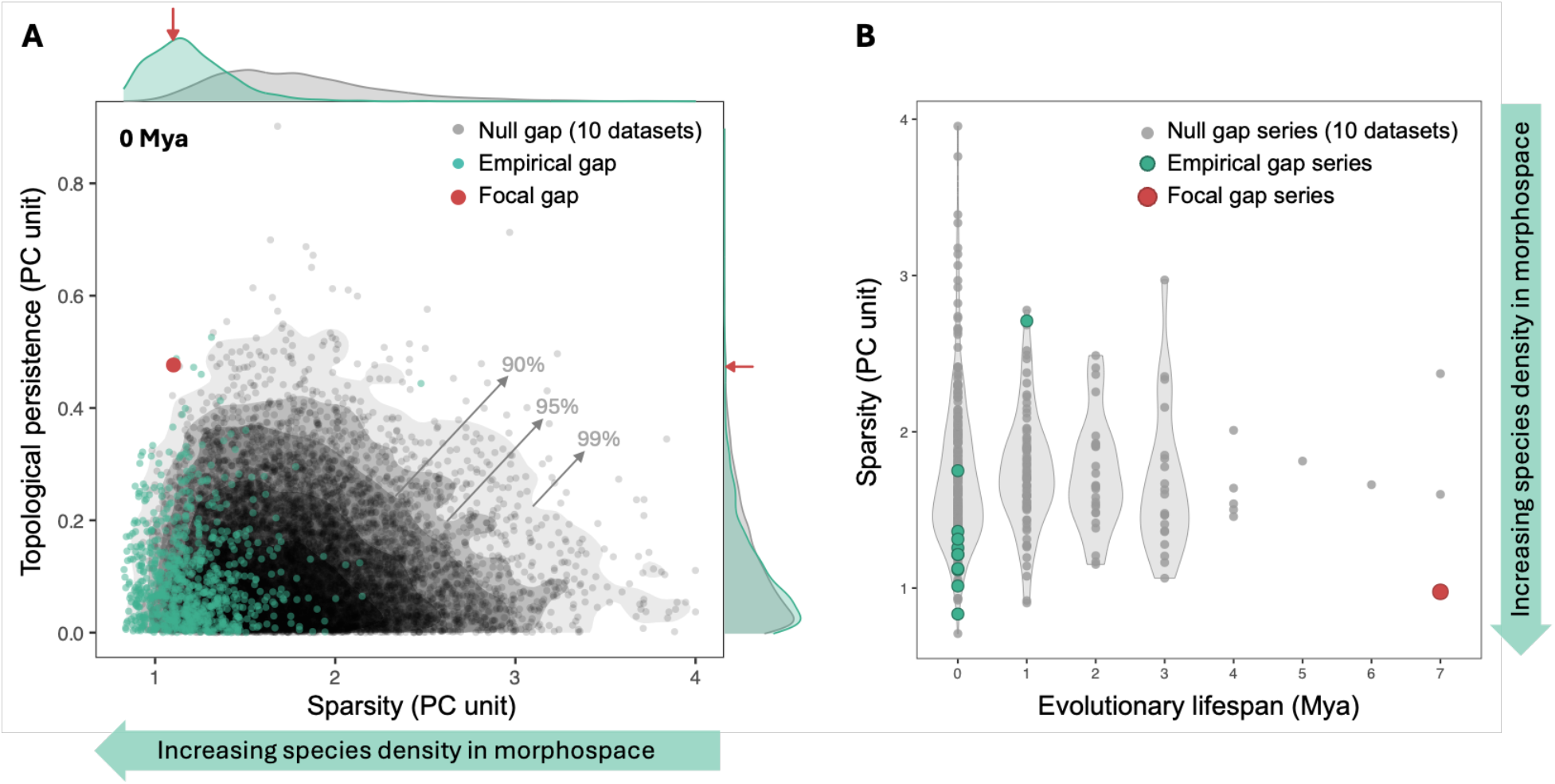
Comparison of the focal morphological gap to gaps detected in null datasets. **A**: Topological persistence versus sparsity for gaps identified at the modern time. Sparsity is defined as the mean distance from the gap centroid to the nearest 5% of data points in the full trait space; lower sparsity indicates denser species distribution in the morphospace. Each point represents a gap from either the empirical dataset (green) or one of 10 null datasets simulating neutral trait evolution (gray). The focal gap is shown in red. Contours indicate the 99%, 95%, and 90% quantiles of the null gap distribution. **B**: Sparsity versus evolutionary lifespan of gap series, where each gap series represents evolutionarily linked gaps detected in similar morphological locations across time slices. The focal gap stands out for its combination of high topological persistence, long evolutionary lifespan (7 million years), and location within a densely occupied region of morphspace.

These differences suggest that the focal gap is unlikely to have emerged under a neutral model of trait diversification, arguing against the neutral diversification hypothesis (**Hypothesis C**).

### 3.3 No evidence for historical occupation or missing data

We next evaluated whether the focal gap might be a remnant of previously existing but now extinct morphologies (**Hypothesis D**). Ancestral trait reconstructions revealed no evidence of occupation or traversal through the gap by ancestral passeroid lineages at any time slice (**Fig. S3**). To complement the limitation that ancestral reconstructions only include ancestors of extant species, we also examined recently extinct species with available measured or imputed morphological trait data (those extinct within the past ∼130,000 years). Among 60 such extinct passeroid species, only three were located near the gap, and none fell within it (**Fig. S4A**). Collectively, these findings provide little support for the extinction hypothesis.

To test whether the gap could be a result of missing data (**Hypothesis E**), we examined the 40 species excluded from our analysis due to incomplete traits. Only one of their reference species (used for trait imputation in the original dataset) fell near the gap (**Fig. S4B**), making it unlikely that the focal gap is caused by missing species clustering in the gap region.

### 3.4 Morphological viability outside the Passeroidea

To evaluate whether the persistent gap reflects an intrinsically infeasible morphology (**Hypothesis B**), we searched for species from other passerine clades that occupy the same region of morphospace. We found multiple non-passeroid species located within the focal gap (**Fig. S5, Table S3**), confirming that this trait combination is biologically viable. Moreover, passeroid species surrounding the gap span multiple families (**Fig. S5, Table S4**), suggesting that the absence of this morphology is not due to a lineage-specific constraint, but instead reflects a broader pattern across the clade. These findings argue against intrinsic constraints as the cause of the gap.

In addition, the non-passeroid gap occupants are geographically widespread (**Fig. 5A**), occupying biogeographic regions where passeroids also occur. This suggests that suitable ecological niches for gap morphologies exist and are accessible to Passeroidea, making the niche absence hypothesis (**Hypothesis F**) unlikely.

**Figure 5.**
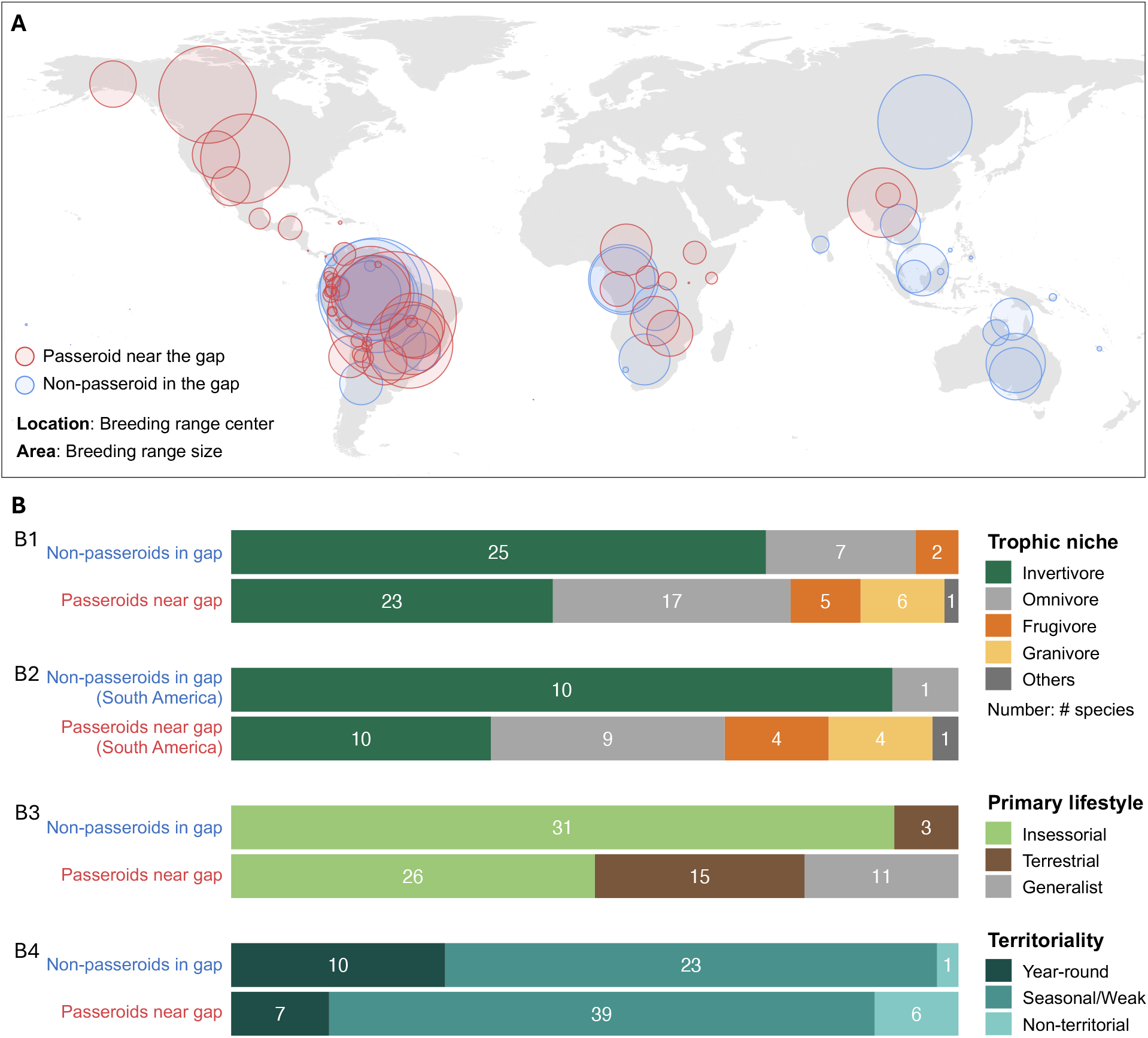
Geographical distribution and ecological traits of non-passeroids in the gap and passeroids near the gap. **A**: Global distribution of breeding range centers for non-passeroids occupying the focal gap (blue) and passeroids near the gap (red). Circle size reflects the true breeding range area (not shape), and location indicates the geographic center of each species’ breeding distribution. **B**: Ecological trait profiles of non-passeroids in the gap and passeroids near the gap. Bar plots show species counts grouped by trophic niche (**B1 & B2**), primary lifestyle (**B3**), and territoriality (**B4**). Trophic niche traits are summarized for all species (**B1**) and separately for species in South America (**B2**), where geographic overlap is most pronounced. The primary lifestyle “insessorial” refers to species that habitually perch above the ground, whether in vegetation or on elevated substrates such as rocks or artificial structures.

### 3.5 Competitive exclusion as the most plausible explanation

That the morphological gap persists within, but not outside, the Passeroidea raises the possibility of competitive exclusion (**Hypothesis G**), and three additional lines of observation provide support for this explanation. First, the non-passeroid gap occupants are predominantly perching insectivores (**Figs. 5B1 & 5B3**) that generally exhibit territorial behavior (**Fig. 5B4**) and are primarily distributed in humid tropical or subtropical forests (**Fig. 5A**). Their narrow niche specialization and capability for resource defense suggest a high potential for competitive behaviors.

Second, phylogenetic and biogeographic evidence indicates that the ancestors of non-passeroid species falling within the morphological gap colonized key habitats earlier than did those of passeroids with similar morphologies. In South America, non-passeroid gap occupants (e.g., Tyrannidae, Tityridae, Furnariidae) arrived during the Oligocene or early Miocene (∼20–30 Mya), whereas passeroids near the gap (e.g., Thraupidae, Icteridae, Passerellidae) arrived later, mostly within the last 3–10 million years following or close to the closure of the Isthmus of Panama (Oliveros *et al*. 2019; Weir *et al*. 2009). A similar pattern is seen in Africa, where the Sylviida clade (e.g. Alaudidae, Pycnonotidae) originated or arrived during the Oligocene (∼25– 30 Mya), preceding the arrival of passeroid lineages like the Ploceidae and Motacillidae, which have Eurasian origins and likely entered Africa only after the early Miocene (Alström *et al*. 2013, 2023; Oliveros *et al*. 2019; Voelker 1999). This historical precedence raises the possibility that early arrivals preempted available ecological niches, limiting opportunities for passeroids to evolve into the same region of trait space.

Third, in South America where the geographic overlap between the two groups is most extensive, the dietary contrast between passeroid and non-passeroid species became more pronounced. Non-passeroid species occupying the gap in morphospace exhibit strong niche specialization, while morphologically similar passeroids display more generalized diets (**Fig. 5B2**), suggesting potential niche displacement of passeroids in response to competition.

Together, these findings support a scenario in which early-arriving, ecologically specialized non-passeroids preempted and defended a distinct ecological niche, thereby constraining passeroid lineages from evolving into that region of morphospace.

## 4. Discussion

### 4.1 Competitive exclusion as the most likely driver of the focal morphological gap

By applying persistent homology to morphological trait data reconstructed through time, we identified a long-standing morphological gap within the Passeroidea morphospace—a region that has remained consistently unoccupied for approximately 7 million years based on the BirdTree phylogeny (Jetz *et al*. 2012). After evaluating seven hypotheses, we ruled out six, leaving competitive exclusion the most plausible explanation. In particular, our findings support a scenario in which early-arriving, ecologically specialized non-passeroids preempted a viable region of morphospace, thereby limiting evolutionary access for passeroid lineages.

The morphology associated with the focal gap is realized by non-passeroid species that are primarily insectivores foraging in vegetation in the tropical system, which has been documented as an ecological strategy linked to high niche specialization and strong interspecific aggression (Robinson & Terborgh 1995; Sherry *et al*. 2020). The idea that early-arriving species can preempt ecological niches is a well-documented phenomenon across diverse taxa (Stroud *et al*. 2024). In the Neotropics, such dynamics have been observed in bird communities in which early colonizing Tyranni dominate in species richness, but later-arriving Passeri exhibit greater functional diversity, likely reflecting niche displacement to avoid competition (Almeida *et al*. 2018). This pattern aligns with our results, where the non-passeroid gap occupants in South America all belong to the Tyranni clade, suggesting that early arrival and ecological specialization may have played a central role in excluding passeroids from evolving into what has for them become a gap in morphospace.

These findings suggest that long-term morphological gaps can arise not just from intrinsic constraints but from historical contingencies in community assembly. Consequently, disruptions to long-standing community structures, such as environmental change and species invasions, may alter competitive dynamics, potentially opening previously inaccessible regions of morphospace or reinforcing new morphological gaps.

### 4.2 Other morphological gaps: neutral and non-neutral drivers

Although the focal gap appears to be a product of long-term competitive dynamics, other gaps identified in the Passeroidea morphospace were generally shorter-lived, with characteristics not particularly deviating from those produced under simulated Brownian motion, suggesting they may have arisen from neutral diversification processes.

Interestingly, compared to null datasets, gaps in the empirical dataset tended to occur in denser regions of morphospace (i.e., those with low sparsity) and were smaller in size (**Figs. 4 and S6**), despite the overall similar variance on the PC axes (**Fig. S7**). This pattern may reflect niche packing, in which closely related species differentiate to coexist in the same environment, leaving smaller gaps. Future research could test this idea by comparing gap patterns between sympatric and allopatric species assemblages, with the expectation that sympatric groups will show lower sparsity and smaller gaps.

### 4.3 Limitations and methodological considerations

Persistent homology is computationally intensive, especially in terms of memory usage for our particular analysis, which limits the ability to explore a broad range of evolutionary models and phylogenetic hypotheses. To maintain minimal assumptions and compatibility with a consensus tree with polytomies, we used ancestral state reconstruction under a Brownian motion model. However, we recognize that this represents only one of many possible evolutionary scenarios and could oversimplify variation in evolutionary rates or selective regimes. Despite these limitations, we believe persistent homology remains a uniquely powerful tool for addressing questions about evolutionary constraints.

The use of a consensus phylogeny containing unresolved polytomies could inflate the spread of ancestral traits, potentially affecting the size or frequency of detected gaps in morphospace. Ideally, such analyses could incorporate numerous fully resolved candidate dichotomous trees to account for phylogenetic uncertainty while maintaining dichotomous tree structure, but the high computational demand currently makes this impractical. Nevertheless, because both the empirical and null datasets were analyzed using the same consensus tree, any biases introduced by tree topology would affect both equally. This makes it unlikely that the distinctive properties of the focal gap are an artifact of tree topology.

The estimated branching time of the Passeroidea has changed considerably over time, from close to 40 million years ago in the BirdTree phylogeny (Jetz *et al*. 2012) to ∼18 million years ago in more recent work (Stiller *et al*. 2024). However, the BirdTree dataset remains the only available global phylogeny for full species-level coverage, and thus was used in our analysis. A more recent divergence time would shift the temporal scale of our results; for example, the focal gap identified as unoccupied for 7 million years may, under updated calibrations, span closer to 3–4 million years. Nonetheless, the central finding that this gap has remained persistently unoccupied throughout the evolutionary history of the Passeroidea remains robust.

Finally, we acknowledge that the ten external morphological traits used in this analysis do not capture the full range of ecological strategies employed by bird species. Furthermore, morphology and ecological niche need not always correspond directly; similar niches may be filled by species with different morphologies, and vice versa. Our framework provides a way to identify interesting patterns observed in trait space and discuss potential mechanisms. Clarifying the specific ecological processes involved, including whether and how interspecific competition acts *in vivo*, will require further investigation.

### 4.4 Broader implications

This study demonstrates the application of persistent homology, a topological data analysis tool, to macroevolutionary questions by examining the structure of unoccupied regions in trait space. By tracing high-dimensional morphological gaps through evolutionary time, we reveal robust features of trait distributions that are often overlooked by conventional dimensionality reduction approaches. This framework also allows us to distinguish short-lived, potentially stochastic gaps from those that are both topologically persistent (stable across scales in trait space) and evolutionary persistent (stable over time).

Although our analysis focuses on the Passeroidea and continuous morphological traits, the approach is broadly applicable across taxonomic groups and trait types, including non-continuous variables (e.g. Nguyen *et al*. 2022). For example, it could be used to identify functional gaps or underutilized climatic niches in plant communities to assess invasion risk, or to detect missing floral trait combinations and their potential associations with pollinator availability. By treating gaps in trait space (i.e., the absence of species with particular trait combinations) as informative biological signals, our approach provides a new perspective on why some organismal forms or strategies do not exist. This novel and generalizable framework integrates trait-based approaches with concepts from evolutionary processes, ecological forces, and biogeographic history to advance our understanding of the drivers and limits of biodiversity.

