## Supplementary Information for "Birds that don’t exist: niche pre-emption as a constraint on morphological evolution in the Passeroidea"

**Figure S1**. Evolutionary trajectories of passeroid morphological traits along the four principal component axes.

**Figure S2**. Bird morphology associated with the focal gap identified in Passeroidea morphospace.

**Figure S3**. Evolutionary trajectories of passeroid morphological traits projected onto PC1 and PC2.

**Figure S4**. Recently extinct and reference species of passeroids projected onto the morphospace defined by the first two PCs of the extant passeroid species included in the analysis.

**Figure S5**. Phylogenetic tree of Passeriformes showing the positions of species near or within the focal morphological gap.

**Figure S6**. Comparison of gap properties between empirical and null datasets.

**Figure S7**. Empirical and simulated null trait distributions in morphospace and their corresponding persistence diagrams.

**Table S1.** Loadings of ten morphological traits onto the principal components for extant Passeroidea species.

**Table S2**. Summary of notable morphological gaps in passeroid morphospace across time slices.

**Table S3**. List of non-passeroid species occupying the focal morphological gap and their ecological traits.

**Table S4**. List of passeroid species near the focal morphological gap and their ecological traits.

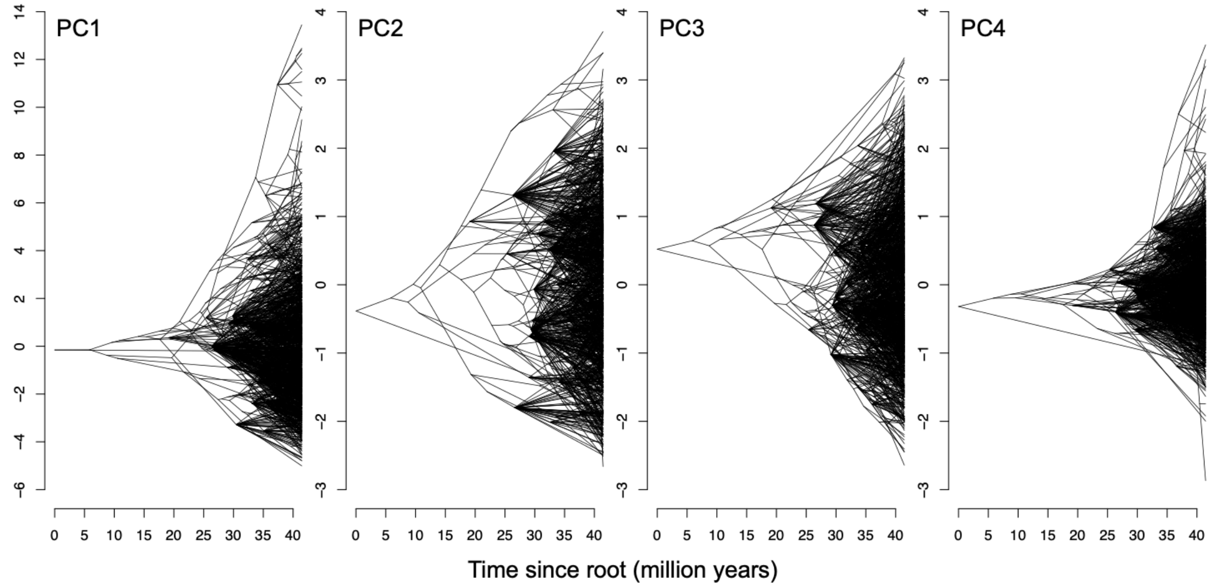


**Figure S1**. Evolutionary trajectories of passeroid morphological traits along the four principal component axes. Time is shown as millions of years since the root of the tree.

**A.** Focal gap centroid trait value

| Trait | Value | Unit |
|---|---|---|
| Beak Length (Culmen) | 17.7 | mm |
| Beak Length (Nares) | 10.6 | mm |
| Beak Width | 4.5 | mm |
| Beak Depth | 5.7 | mm |
| Wing Length | 83.1 | mm |
| Secondary Length | 67.7 | mm |
| Hand-Wing Index | 18.2 | - |
| Tarsus Length | 24.1 | mm |
| Tail Length | 71.9 | mm |
| Body Mass | 27.7 | gram |

**B.** Non-passeroid species within the focal gap

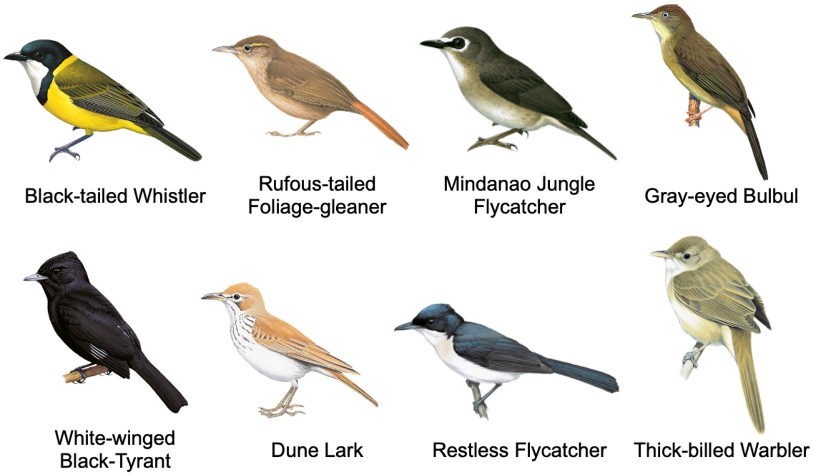


**Figure S2**. Bird morphology associated with the focal gap identified in Passeroidea morphospace. **A**: Morphological trait values corresponding to the centroid of the focal gap. **B**: Illustrations of example non-passeroid species that occupy the focal gap region of trait space, sourced from Birds of the World, Cornell Lab of Ornithology.

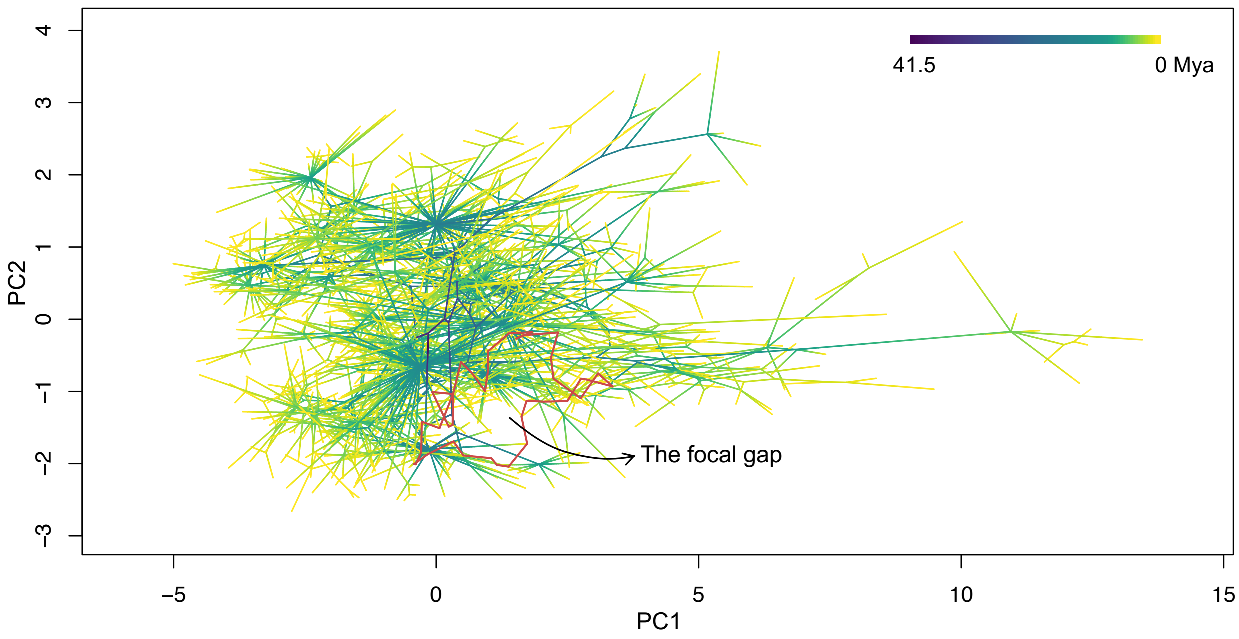


**Figure S3**. Evolutionary trajectories of passeroid morphological traits projected onto PC1 and PC2. Lines trace reconstructed trait changes along the phylogeny, based on ancestral state estimation under a Brownian motion model. Color represents evolutionary time, with darker lines indicating earlier time points (further in the past) and lighter lines indicating more recent events. Red loop marks the focal gap. Note that this is a 2D projection of a 4D trait space; while some trajectories appear to pass through the focal gap in this projection, they do not in the full space. The persistence of the focal gap over evolutionary time (**Fig. 3**) further supports this interpretation.

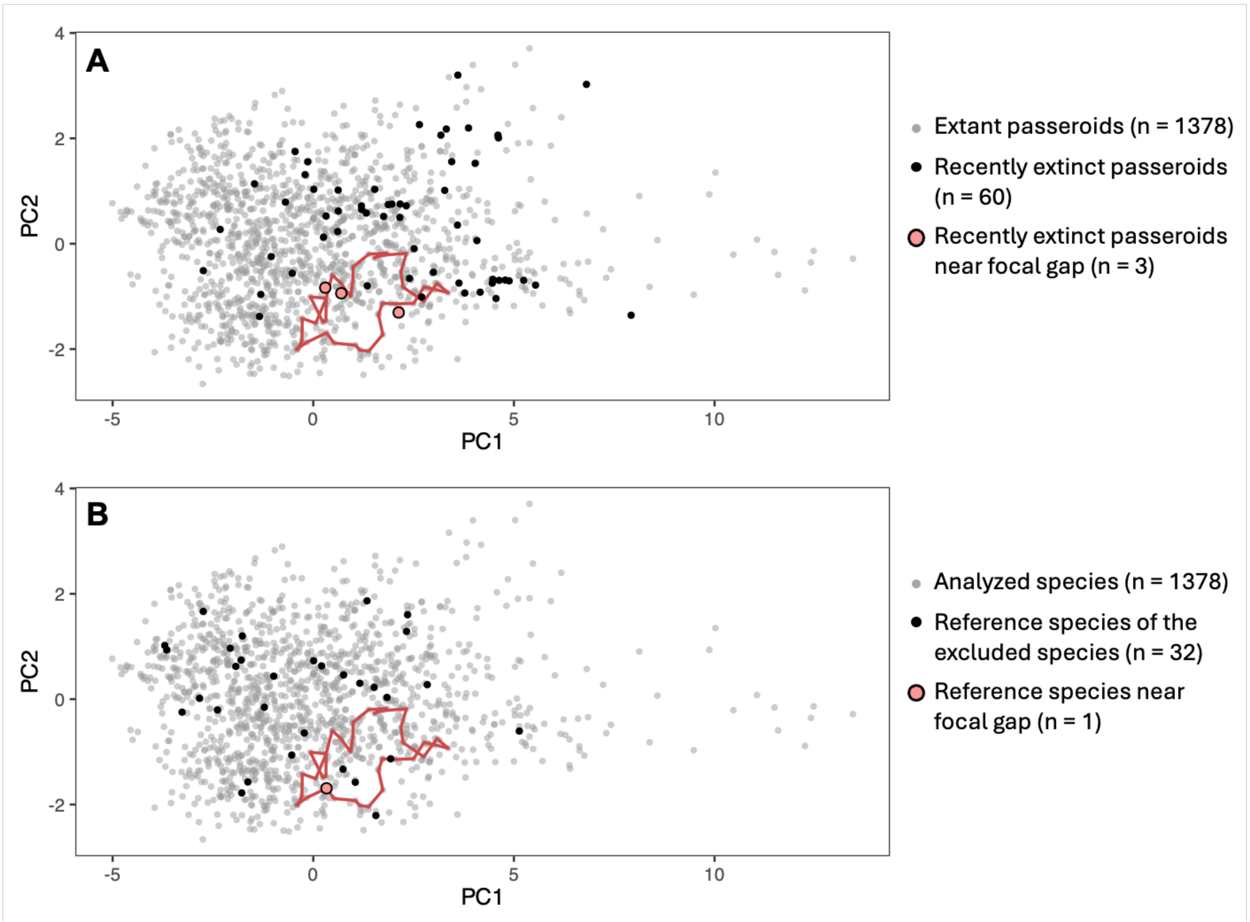


**Figure S4**. Recently extinct and reference species of passeroids projected onto the morphospace defined by the first two PCs of the extant passeroid species included in the analysis. **A**: Positions of 60 recently extinct passeroid species (black points) and extant passeroids (gray points). Three recently extinct species near the gap are shown in pink. **B**: Positions of 32 reference species (black points) used for trait imputation (based on the original dataset; Tobias et al., 2022) for the excluded passeroid species, and other passeroid species included in the analysis (gray points). One reference species located near the gap is shown in pink. The red outline in both panels indicates the focal morphological gap. These plots are 2D projections of a 4D trait space; while some species appear to fall within the focal gap in this projection, they do not in the full space.

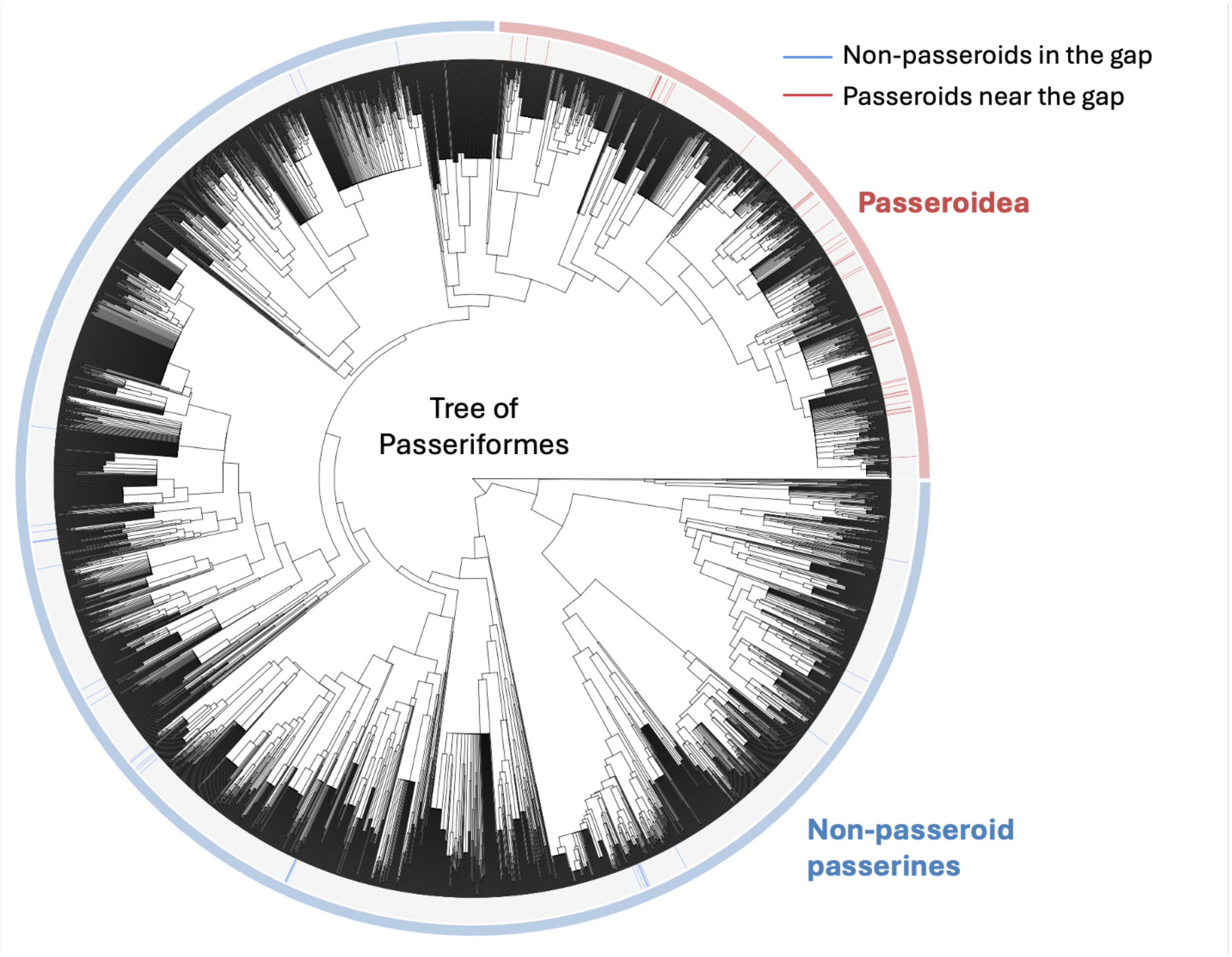


**Figure S5**. Phylogenetic tree of Passeriformes showing the positions of species near or within the focal morphological gap. Species near the gap within the Passeroidea (n = 52) are marked in red, and non-passeroid species occupying the gap (n = 34) are marked in blue. This distribution indicates that species associated with the focal gap morphology are phylogenetically dispersed across multiple lineages.

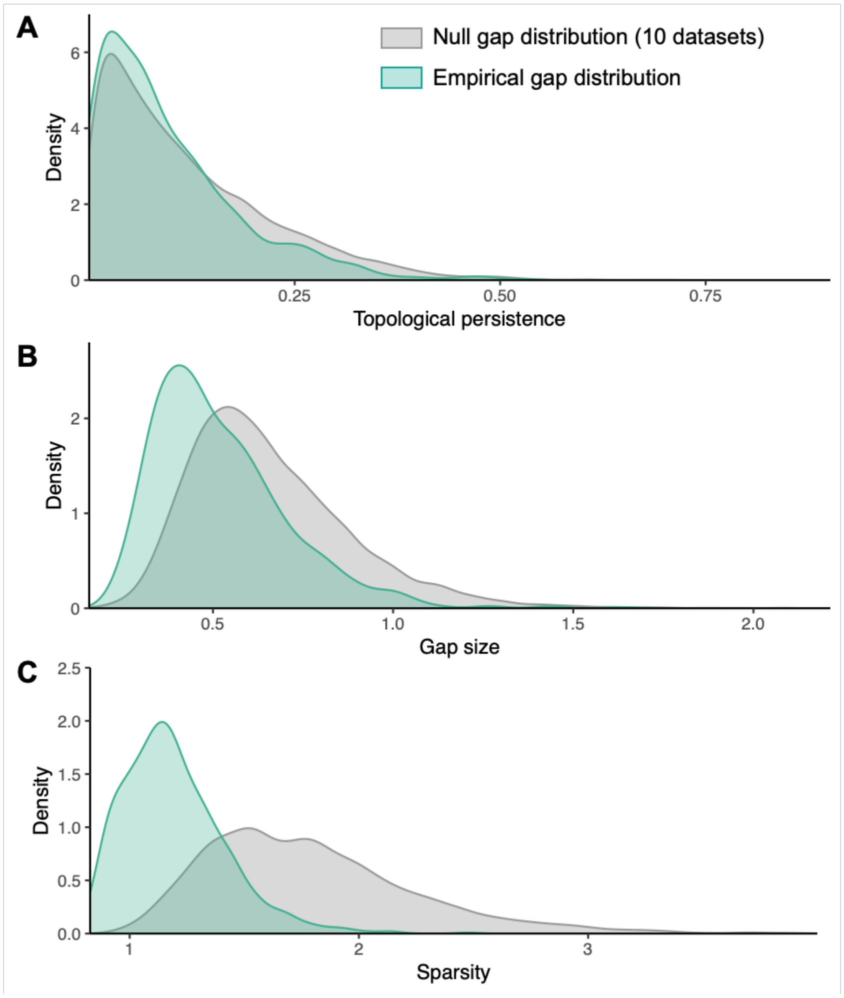


**Figure S6**. Comparison of gap properties between empirical and null datasets. Distributions of topological persistence (**A**), size (**B**), and sparsity (**C**) are shown for gaps detected in empirical dataset (green) and in null datasets simulated under a Brownian motion model (gray). Compared to the null expectation, empirical gaps tend to be smaller and located in denser regions in the morphospace, suggesting they may reflect non-random evolutionary process.

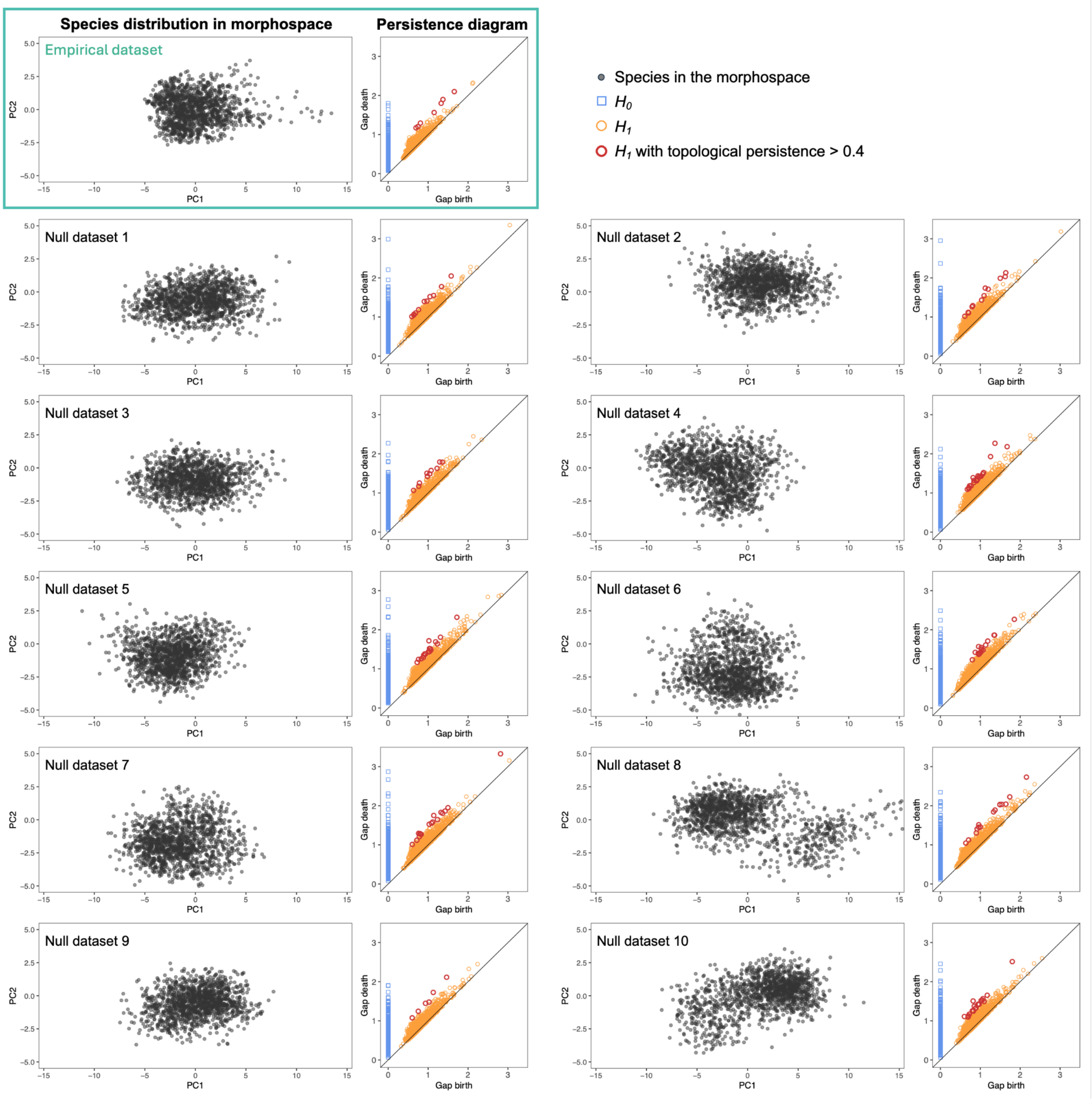


**Figure S7**. Empirical and simulated null trait distributions in morphospace and their corresponding persistence diagrams. The top-left panel shows the empirical dataset; the remaining panels show the 10 null datasets simulated under a Brownian motion model using the same parameters, all at the modern time slice. In each panel, the scatter plot (left) shows species projected onto the PC1 x  PC2 plane. Corresponding persistence diagrams (right) show $H_0$ features (connected components, blue squares) and $H_1$ features (loops, orange circles), with notable loop structures (Gap birth - Gap death, or topological persistence, larger than 0.4) outlined in red. One $H_0$ feature with a particularly high gap death value is excluded from each persistence diagram for visualization clarity. Trait distributions in the null datasets exhibit similar variance to the empirical dataset, as expected under matched simulation parameters.

**Table S1.** Loadings of ten morphological traits onto the principal components for extant Passeroidea species.

| | PC1 | PC2 | PC3 | PC4 | PC5 | PC6 | PC7 | PC8 | PC9 | PC10 |
|---|---|---|---|---|---|---|---|---|---|---|
| Beak Length (Culmen) | 0.35 | -0.12 | -0.03 | -0.52 | 0.13 | -0.09 | 0.10 | 0.02 | 0.74 | 0.00 |
| Beak Length (Nares) | 0.35 | 0.04 | 0.01 | -0.59 | 0.36 | 0.00 | -0.08 | -0.01 | -0.63 | 0.01 |
| Beak Width | 0.25 | 0.60 | 0.24 | 0.11 | 0.00 | 0.17 | 0.57 | -0.40 | 0.02 | 0.00 |
| Beak Depth | 0.27 | 0.54 | 0.27 | 0.12 | -0.02 | 0.11 | -0.67 | 0.27 | 0.15 | -0.01 |
| Tarsus Length | 0.33 | -0.37 | 0.07 | -0.04 | -0.46 | 0.70 | -0.10 | -0.17 | -0.06 | 0.00 |
| Wing Length | 0.36 | -0.04 | -0.30 | 0.20 | -0.11 | -0.32 | -0.15 | -0.32 | -0.04 | -0.71 |
| Secondary Length | 0.37 | -0.15 | -0.03 | 0.22 | -0.12 | -0.42 | -0.16 | -0.34 | -0.04 | 0.67 |
| Hand-wing index (HWI) | 0.01 | 0.32 | -0.88 | -0.01 | 0.00 | 0.27 | -0.02 | 0.04 | 0.04 | 0.22 |
| Tail Length | 0.32 | -0.27 | -0.01 | 0.51 | 0.67 | 0.25 | 0.11 | 0.19 | 0.05 | 0.00 |
| Body Mass | 0.37 | 0.01 | -0.04 | 0.07 | -0.40 | -0.21 | 0.38 | 0.69 | -0.15 | 0.00 |
| Variance explained (%) | 66.11 | 13.29 | 11.01 | 4.60 | 1.93 | 1.36 | 0.72 | 0.64 | 0.33 | 0.01 |
| Cumulative variance (%) | 66.11 | 79.40 | 90.42 | 95.01 | 96.94 | 98.30 | 99.02 | 99.66 | 99.99 | 100.00 |

**Table S2**. Summary of notable morphological gaps in passeroid morphospace across time slices. Each row corresponds to a gap detected at a specific time slice (Mya = million years ago) that exceeds the topological persistence threshold (0.4). Columns show the number of vertices forming the gap (i.e., the number of species defining the enclosing loop), evolutionary lifespan (in million years), gap size (mean distance from centroid to vertices), and local sparsity (mean distance to nearest 5% of species). A gap series links morphological gaps across time, with the color of group IDs matching those shown in **Figure 2**. Group 1 corresponds to the focal gap.

| Mya | Number of vertices | Topological persistence | Gap size | Location sparsity | Gap series index |
|---|---|---|---|---|---|
| 0 | 38 | 0.48 | 1.27 | 1.10 | **1** |
| 1 | 39 | 0.57 | 1.39 | 1.08 | **1** |
| 2 | 35 | 0.49 | 1.45 | 1.02 | **1** |
| 3 | 36 | 0.53 | 1.49 | 1.02 | **1** |
| 4 | 29 | 0.64 | 1.46 | 0.95 | **1** |
| 5 | 38 | 0.70 | 1.54 | 0.81 | **1** |
| 6 | 22 | 0.54 | 1.36 | 0.82 | **1** |
| 7 | 18 | 0.44 | 1.25 | 1.00 | **1** |
| 0 | 13 | 0.46 | 0.87 | 1.26 | **2** |
| 0 | 21 | 0.49 | 1.14 | 1.12 | **3** |
| 0 | 9 | 0.41 | 0.98 | 1.36 | **4** |
| 0 | 6 | 0.47 | 1.00 | 1.21 | **5** |
| 0 | 5 | 0.53 | 1.03 | 1.31 | **6** |
| 0 | 10 | 0.44 | 1.42 | 2.48 | **7** |
| 1 | 6 | 0.41 | 1.13 | 2.94 | **7** |
| 1 | 17 | 0.41 | 1.10 | 1.01 | **8** |
| 1 | 8 | 0.52 | 1.20 | 1.22 | **9** |
| 3 | 16 | 0.42 | 0.91 | 1.75 | **10** |
| 5 | 19 | 0.46 | 0.94 | 1.12 | **11** |
| 6 | 31 | 0.45 | 1.07 | 0.83 | **12** |

**Table S3**. List of non-passeroid species occupying the focal morphological gap and their ecological traits.

| Species | Family | Order | Trophic niche | Primary lifestyle | Habitat | Territoriality | Zoological realm |
|---|---|---|---|---|---|---|---|
| *Jynx ruficollis* | Picidae | Piciformes | Invertivore | Insessorial | Woodland | Year-round | Afrotropical |
| *Deconychura stictolaema* | Furnariidae | Passeriformes | Invertivore | Insessorial | Forest | Year-round | Neotropical |
| *Anabacerthia striaticollis* | Furnariidae | Passeriformes | Invertivore | Insessorial | Forest | Seasonal/Weak | Neotropical |
| *Philydor atricapillus* | Furnariidae | Passeriformes | Invertivore | Insessorial | Forest | Seasonal/Weak | Neotropical |
| *Philydor erythrocercum* | Furnariidae | Passeriformes | Invertivore | Insessorial | Forest | Seasonal/Weak | Neotropical |
| *Philydor fuscipenne* | Furnariidae | Passeriformes | Invertivore | Insessorial | Forest | Year-round | Neotropical |
| *Philydor ruficaudatum* | Furnariidae | Passeriformes | Invertivore | Insessorial | Forest | Seasonal/Weak | Neotropical |
| *Syndactyla roraimae* | Furnariidae | Passeriformes | Invertivore | Insessorial | Forest | Year-round | Neotropical |
| *Schiffornis turdina* | Tityridae | Passeriformes | Omnivore | Insessorial | Forest | None | Neotropical |
| *Casiornis rufus* | Tyrannidae | Passeriformes | Invertivore | Insessorial | Woodland | Seasonal/Weak | Neotropical |
| *Knipolegus aterrimus* | Tyrannidae | Passeriformes | Invertivore | Insessorial | Shrubland | Seasonal/Weak | Neotropical |
| *Myiozetetes similis* | Tyrannidae | Passeriformes | Invertivore | Insessorial | Shrubland | Seasonal/Weak | Neotropical |
| *Pachycephala caledonica* | Pachycephalidae | Passeriformes | Invertivore | Insessorial | Forest | Year-round | Oceanian |
| *Pachycephala flavifrons* | Pachycephalidae | Passeriformes | Invertivore | Insessorial | Forest | Year-round | Oceanian |
| *Pachycephala melanura* | Pachycephalidae | Passeriformes | Invertivore | Insessorial | Woodland | Year-round | Oceanian/Australian |
| *Pachycephala pectoralis* | Pachycephalidae | Passeriformes | Invertivore | Insessorial | Forest | Year-round | Australian |
| *Monarcha verticalis* | Monarchidae | Passeriformes | Invertivore | Insessorial | Forest | Seasonal/Weak | Oceanian |
| *Myiagra alecto* | Monarchidae | Passeriformes | Invertivore | Insessorial | Riverine | Seasonal/Weak | Oceanian |
| *Myiagra inquieta* | Monarchidae | Passeriformes | Invertivore | Insessorial | Woodland | Seasonal/Weak | Oceanian/Australian |
| *Pomarea dimidiata* | Monarchidae | Passeriformes | Invertivore | Insessorial | Forest | Year-round | Oceanian |
| *Pomarea iphis* | Monarchidae | Passeriformes | Invertivore | Insessorial | Forest | Seasonal/Weak | Oceanian |
| *Certhilauda barlowi* | Alaudidae | Passeriformes | Omnivore | Terrestrial | Desert | Seasonal/Weak | Afrotropical |
| *Certhilauda erythrochlamys* | Alaudidae | Passeriformes | Omnivore | Terrestrial | Desert | Seasonal/Weak | Afrotropical |
| *Mirafra apiata* | Alaudidae | Passeriformes | Invertivore | Terrestrial | Grassland | Seasonal/Weak | Afrotropical |
| *Andropadus gracilirostris* | Pycnonotidae | Passeriformes | Frugivore | Insessorial | Forest | Seasonal/Weak | Afrotropical |
| *Iole indica* | Pycnonotidae | Passeriformes | Omnivore | Insessorial | Forest | Seasonal/Weak | Oriental |
| *Iole propinqua* | Pycnonotidae | Passeriformes | Omnivore | Insessorial | Forest | Seasonal/Weak | Oriental |
| *Ixos palawanensis* | Pycnonotidae | Passeriformes | Omnivore | Insessorial | Forest | Seasonal/Weak | Oriental |
| *Pycnonotus eutilotus* | Pycnonotidae | Passeriformes | Frugivore | Insessorial | Forest | Seasonal/Weak | Oriental |
| *Pycnonotus goiavier* | Pycnonotidae | Passeriformes | Omnivore | Insessorial | Shrubland | Seasonal/Weak | Oriental |
| *Acrocephalus aedon* | Sylviidae | Passeriformes | Invertivore | Insessorial | Shrubland | Seasonal/Weak | Palearctic/Oriental |
| *Fraseria ocreata* | Muscicapidae | Passeriformes | Invertivore | Insessorial | Forest | Year-round | Afrotropical |
| *Rhinomyias goodfellowi* | Muscicapidae | Passeriformes | Invertivore | Insessorial | Forest | Seasonal/Weak | Oriental |
| *Rhinomyias gularis* | Muscicapidae | Passeriformes | Invertivore | Insessorial | Forest | Seasonal/Weak | Oriental |

Note: "Insessorial" refers to species that habitually perch above the ground, whether in vegetation or on elevated substrates such as rocks or artificial structures.

**Table S4**. List of passeroid species near the focal morphological gap and their ecological traits.

| Species | Family | Order | Trophic niche | Primary lifestyle | Habitat | Territory | Zoological realm |
|---|---|---|---|---|---|---|---|
| *Plocepasser superciliosus* | Ploceidae | Passeriformes | Granivore | Terrestrial | Woodland | Year-round | Afrotropical |
| *Ploceus insignis* | Ploceidae | Passeriformes | Invertivore | Insessorial | Forest | Seasonal/Weak | Afrotropical |
| *Malimbus coronatus* | Ploceidae | Passeriformes | Invertivore | Insessorial | Forest | Year-round | Afrotropical |
| *Anthus sylvanus* | Motacillidae | Passeriformes | Invertivore | Terrestrial | Grassland | Seasonal/Weak | Afrotropical |
| *Anthus lineiventris* | Motacillidae | Passeriformes | Invertivore | Terrestrial | Grassland | Seasonal/Weak | Afrotropical |
| *Anthus similis* | Motacillidae | Passeriformes | Invertivore | Terrestrial | Grassland | Seasonal/Weak | Afrotropical |
| *Anthus melindae* | Motacillidae | Passeriformes | Invertivore | Terrestrial | Grassland | Seasonal/Weak | Afrotropical |
| *Anthus pallidiventris* | Motacillidae | Passeriformes | Invertivore | Terrestrial | Grassland | Seasonal/Weak | Afrotropical |
| *Macronyx flavicollis* | Motacillidae | Passeriformes | Invertivore | Terrestrial | Grassland | Seasonal/Weak | Afrotropical |
| *Macronyx ameliae* | Motacillidae | Passeriformes | Invertivore | Terrestrial | Grassland | Seasonal/Weak | Afrotropical |
| *Macronyx sharpei* | Motacillidae | Passeriformes | Invertivore | Terrestrial | Grassland | Seasonal/Weak | Afrotropical |
| *Melophus lathami* | Emberizidae | Passeriformes | Granivore | Terrestrial | Grassland | Seasonal/Weak | Oriental |
| *Chlorospingus inornatus* | Passerellidae | Passeriformes | Frugivore | Insessorial | Forest | None | Panamanian |
| *Chlorospingus flavigularis* | Passerellidae | Passeriformes | Frugivore | Insessorial | Forest | None | Neotropical |
| *Pipilo chlorurus* | Passerellidae | Passeriformes | Omnivore | Generalist | Shrubland | Seasonal/Weak | Nearctic |
| *Atlapetes melanopsis* | Passerellidae | Passeriformes | Omnivore | Generalist | Shrubland | Year-round | Neotropical |
| *Atlapetes fulviceps* | Passerellidae | Passeriformes | Omnivore | Insessorial | Forest | Seasonal/Weak | Neotropical |
| *Junco vulcani* | Passerellidae | Passeriformes | Omnivore | Generalist | Shrubland | Seasonal/Weak | Panamanian |
| *Zonotrichia leucophrys* | Passerellidae | Passeriformes | Omnivore | Generalist | Shrubland | Seasonal/Weak | Nearctic |
| *Zonotrichia atricapilla* | Passerellidae | Passeriformes | Omnivore | Generalist | Shrubland | Seasonal/Weak | Nearctic |
| *Icteria virens* | Icteridae | Passeriformes | Invertivore | Insessorial | Forest | Seasonal/Weak | Nearctic/Panamanian |
| *Icterus cucullatus* | Icteridae | Passeriformes | Omnivore | Insessorial | Woodland | Seasonal/Weak | Nearctic/Panamanian |
| *Icterus prosthemelas* | Icteridae | Passeriformes | Omnivore | Insessorial | Forest | Year-round | Panamanian |
| *Icterus auricapillus* | Icteridae | Passeriformes | Omnivore | Insessorial | Woodland | Seasonal/Weak | Neotropical |
| *Icterus cayanensis* | Icteridae | Passeriformes | Invertivore | Insessorial | Forest | Seasonal/Weak | Neotropical |
| *Chrysomus ruficapillus* | Icteridae | Passeriformes | Granivore | Insessorial | Wetland | None | Neotropical |
| *Euthlypis lachrymosa* | Parulidae | Passeriformes | Invertivore | Generalist | Woodland | Seasonal/Weak | Nearctic/Panamanian |
| *Creurgops verticalis* | Thraupidae | Passeriformes | Invertivore | Insessorial | Forest | None | Neotropical |
| *Trichothraupis melanops* | Thraupidae | Passeriformes | Invertivore | Insessorial | Forest | Seasonal/Weak | Neotropical |
| *Tachyphonus surinamus* | Thraupidae | Passeriformes | Omnivore | Insessorial | Forest | None | Neotropical |
| *Cnemoscopus rubrirostris* | Thraupidae | Passeriformes | Invertivore | Insessorial | Forest | Seasonal/Weak | Neotropical |
| *Poospiza boliviana* | Thraupidae | Passeriformes | Omnivore | Generalist | Shrubland | Seasonal/Weak | Neotropical |
| *Hemispingus atropileus* | Thraupidae | Passeriformes | Invertivore | Insessorial | Forest | Seasonal/Weak | Neotropical |
| *Hemispingus auricularis* | Thraupidae | Passeriformes | Invertivore | Insessorial | Forest | Seasonal/Weak | Neotropical |
| *Cypsnagra hirundinacea* | Thraupidae | Passeriformes | Invertivore | Insessorial | Grassland | Year-round | Neotropical |
| *Poospiza hypochondria* | Thraupidae | Passeriformes | Invertivore | Insessorial | Shrubland | Seasonal/Weak | Neotropical |
| *Nesospiza acunhae* | Thraupidae | Passeriformes | Omnivore | Generalist | Shrubland | Seasonal/Weak | Atlantic Island |
| *Nesospiza questi* | Thraupidae | Passeriformes | Omnivore | Generalist | Shrubland | Seasonal/Weak | Atlantic Island |
| *Phrygilus unicolor* | Thraupidae | Passeriformes | Granivore | Terrestrial | Grassland | Seasonal/Weak | Neotropical |
| *Phrygilus erythronotus* | Thraupidae | Passeriformes | Granivore | Terrestrial | Grassland | Seasonal/Weak | Neotropical |
| *Phrygilus dorsalis* | Thraupidae | Passeriformes | Granivore | Terrestrial | Grassland | Seasonal/Weak | Neotropical |
| *Diglossa major* | Thraupidae | Passeriformes | Nectarivore | Insessorial | Forest | Year-round | Neotropical |
| *Iridosornis reinhardti* | Thraupidae | Passeriformes | Frugivore | Insessorial | Forest | Seasonal/Weak | Neotropical |
| *Iridosornis rufivertex* | Thraupidae | Passeriformes | Omnivore | Insessorial | Forest | Seasonal/Weak | Neotropical |
| *Delothraupis castaneoventris* | Thraupidae | Passeriformes | Omnivore | Insessorial | Forest | Seasonal/Weak | Neotropical |
| *Thraupis cyanocephala* | Thraupidae | Passeriformes | Frugivore | Insessorial | Forest | None | Neotropical |
| *Anisognathus lacrymosus* | Thraupidae | Passeriformes | Frugivore | Insessorial | Forest | Seasonal/Weak | Neotropical |
| *Neothraupis fasciata* | Thraupidae | Passeriformes | Invertivore | Generalist | Woodland | Year-round | Neotropical |
| *Paroaria baeri* | Thraupidae | Passeriformes | Invertivore | Generalist | Shrubland | Seasonal/Weak | Neotropical |
| *Paroaria capitata* | Thraupidae | Passeriformes | Omnivore | Terrestrial | Shrubland | Seasonal/Weak | Neotropical |
| *Paroaria gularis* | Thraupidae | Passeriformes | Omnivore | Terrestrial | Shrubland | Seasonal/Weak | Neotropical |
| *Phaenicophilus poliocephalus* | Phaenicophilidae | Passeriformes | Invertivore | Insessorial | Forest | Seasonal/Weak | Oriental |

Note: "Insessorial" refers to species that habitually perch above the ground, whether in vegetation or on elevated substrates such as rocks or artificial structures.
